# *Fam72a* controls the balance between error-prone and error-free DNA repair during antibody diversification

**DOI:** 10.1101/2020.12.22.424012

**Authors:** Mélanie Rogier, Jacques Moritz, Isabelle Robert, Chloé Lescale, Vincent Heyer, Anne-Sophie Thomas-Claudepierre, Arthur Abello, Ludovic Deriano, Bernardo Reina-San-Martin

## Abstract

Efficient humoral responses rely on DNA damage, mutagenesis and error-prone DNA repair. B cell receptor diversification through somatic hypermutation (SHM) and class switch recombination (CSR) are initiated by cytidine deamination in DNA mediated by activation induced cytidine deaminase (AID)^1^ and by the subsequent excision of the resulting uracils by Uracil DNA glycosylase (UNG) and by mismatch repair (MMR) proteins^2–4^. Although uracils arising in DNA are faithfully repaired^2–7^, it is not known how these pathways are co-opted to generate mutations and double stranded DNA breaks (DSBs) in the context of SHM and CSR^2,4,8^. Here we have performed a genome-wide CRISPR/Cas9 knockout screen for genes involved in CSR. The screen identified FAM72A, a protein that interacts with the nuclear isoform of UNG (UNG2)^9^ and that is overexpressed in several cancers^9^. We show that the FAM72A-UNG2 interaction controls the protein levels of UNG2 and that CSR is defective in *Fam72a^−/−^* B cells due to the specific upregulation of UNG2. Moreover, we show that in *Fam72a^−/−^* B cells SHM is reduced by 5-fold and that upregulation of UNG2 results in a skewed mutation pattern. Our results are consistent with a model in which FAM72A interacts with UNG2 to control its physiological level by triggering its degradation. Consequently, deficiency in *Fam72a* leads to supraphysiological levels of UNG2 and enhanced uracil excision, shifting the balance from error-prone to error-free DNA repair. Our findings have potential implications for tumorigenesis, as *Fam72a* overexpression would lead to reduced UNG2 levels, shifting the balance toward mutagenic DNA repair and rendering cells more prone to acquire mutations.

---

During immune responses, mammalian B cells undergo somatic hypermutation (SHM) and class switch recombination (CSR) to diversify the B cell receptor repertoire and mount efficient and adapted humoral responses^2,3^. SHM modifies antibody affinity for the triggering antigen by introducing point mutations at the variable region of both heavy (IgH) and light (IgL) chains, thereby allowing affinity maturation and the subsequent generation of high-affinity antibodies^2,3^. CSR modulates antibody effector functions by replacing the isotype expressed from IgM/IgD to IgG, IgA or IgE^10^. This is achieved by a deletional recombination event that takes place at the IgH locus between a donor (Sμ) and an acceptor switch region (Sx)^10^ that brings into proximity the V region and the exons encoding for a new constant region, thus allowing the expression of an antibody with the same specificity but with a different isotype^10^. Mechanistically, both SHM and CSR are triggered by activation induced cytidine deaminase (AID), an enzyme which deaminates cytosines into uracils in DNA^11^. AID-generated U:G mismatches are processed mainly by the base-excision repair (BER) enzyme uracil-DNA-glycosylase (UNG)^4,12^ but also by mismatch repair (MMR)^13^ proteins to introduce mutations or double-stranded DNA breaks (DSBs) during SHM and CSR, respectively^2–4^. During SHM, DNA replication over U:G mismatches generates transition mutations at G:C base pairs. Uracil excision by UNG, followed by replication over abasic sites introduces both transition and transversion mutations at C:G base pairs. In addition, single stranded DNA surrounding the U:G mismatch or abasic site can be excised by both MMR and BER to introduce transition and transversion mutations at A:T base pairs. During CSR, cytidine deamination and DNA cleavage on opposite strands generate DSBs^10^, which are a necessary intermediates for recombination^3^. While uracils in DNA are usually faithfully repaired^2–7^, the mechanisms by which AID-induced uracils are instead processed in an error-prone way in the context of antibody diversification through SHM and CSR is largely unknown^2–4^.

To get insight into the molecular mechanisms controlling CSR, we conducted a genome-wide CRISPR/Cas9 knockout screen (**Fig. 1A**). The screen was performed with the second-generation mouse Brie (mBrie)^14^ gRNA library and using the IgM^+^ murine B cell line CH12, which can be induced to express endogenous levels of AID and undergo CSR from IgM to IgA very efficiently *in vitro* when cultured with TGF-β, IL-4 and an anti-CD40 antibody^15^. To conduct the screen, we established a CH12 cell line stably expressing Cas9 (CH12^Cas9^; **Fig. S1A**) and sub-cloned the gRNA library into a retroviral vector (pMX-mBrie; **Fig. S1B**), which efficiently transduces CH12 cells. CH12^Cas9^ cells were transduced with the pMX-mBrie gRNA library and selected with puromycin for two weeks to allow for the Cas9-mediated generation of knockouts. Cells were then stimulated to undergo CSR for 72h. Cells which failed to undergo CSR (which remained IgM+) or that succeed (which became IgA+) were sorted using magnetic beads (**Fig. 1A and S1C**). Genes, whose loss-of-function affect the efficiency of CSR, were identified by looking at gRNA enrichment in the IgM+ versus IgA+ populations through deep sequencing^16^ (**Fig. 1A**). The screen identified 654 candidate genes with a significant (p<0.05) gRNA enrichment and having more than 2 effective gRNAs (**Fig. 1B**). The screen was successful as it identified genes known to be required for CSR (*Aicda, Ung, Trp53bp1*) together with transcription factors driving AID expression (*Batf, Pten, Irf4*), Non-homologous end joining factors (*Xrcc5, Fam35a*), mismatch repair proteins (*Msh2, Msh6, Pms1, Pms2, Mlh1*), members of the TGF-β, IL-4 and CD40 signaling pathways (*Tfgbr1, Tgfbr2, Il4ra, Il2rg, CD40*) and other genes known to be implicated in CSR (*Mediator, Trim28, Zmynd8*, etc.). Consistent with this, gene ontology analysis revealed a significant enrichment in pathways that are relevant for CSR and SHM (**Fig. 1C**). Importantly, the screen revealed a significant number of genes with no known function in CSR, including *Fam72a* (**Fig. 1B**).

**Figure 1.**
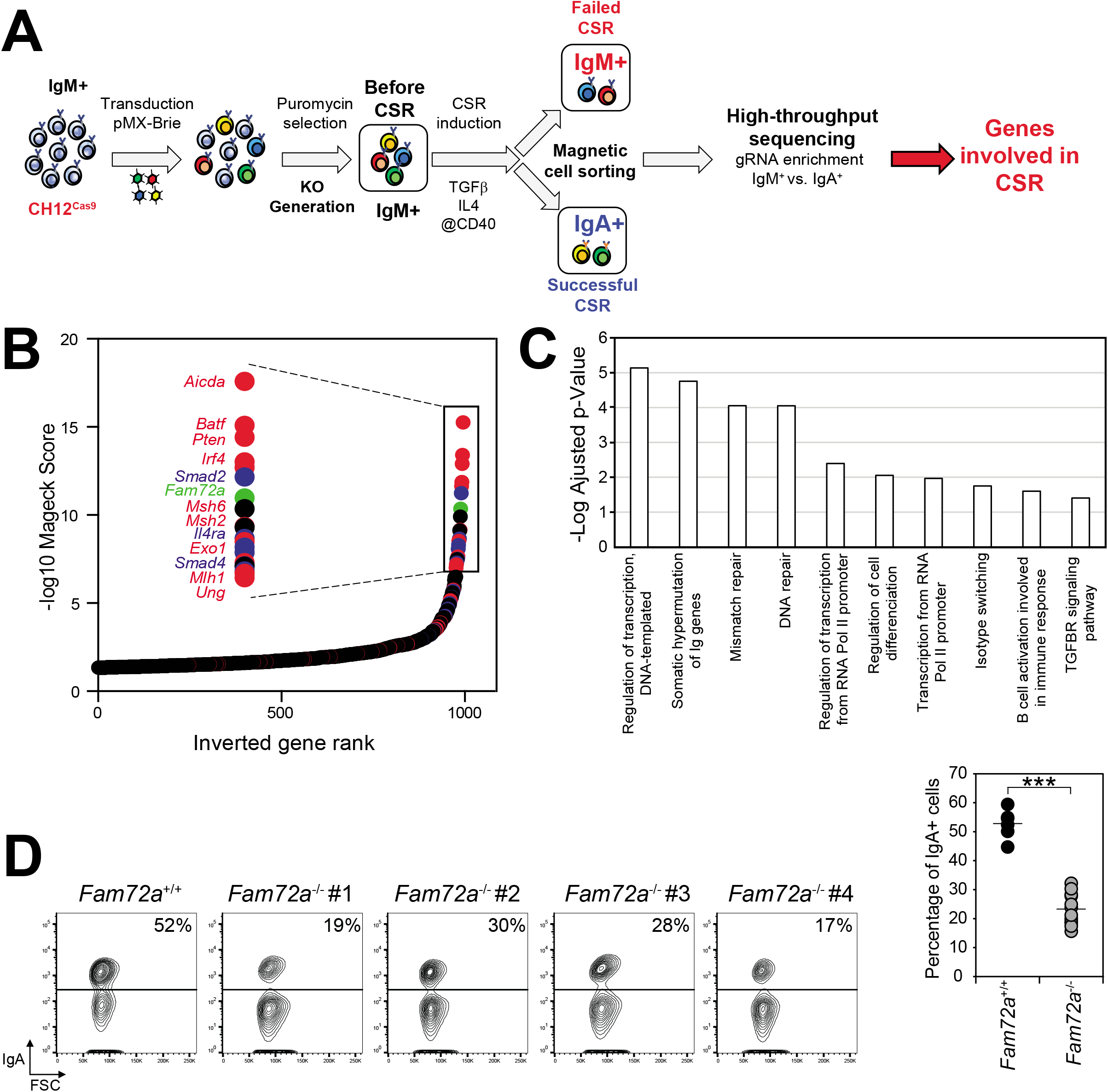
A genome-wide CRISPR/Cas9 knockout screen for genes involved in CSR identifies Fam72a. **(A)** Schematic overview of the CRISPR/Cas9 screen. **(B)** Graph depicting the gene significance score of the ranking (-log10 Score Mageck) calculated with the MAGeCK algorithm, plotted against the logarithmic value to the base 2 of the ratio of IgM over IgA read counts (LFC) for each gene represented by a dot. Genes known to be involved in CSR are depicted in red, TGF-β, IL-4, CD40 and BCR signaling pathways are depicted in blue. Genes with no known function in CSR are in black. **(C)** Gene ontology analysis of genes identified in the screen. **(D)** Flow cytometry analysis of IgA expression in *Fam72a^+/+^* and four independent *Fam72a^−/−^* CH12 cell clones cultured for 3 days with TGF-β, anti-CD40 antibody and IL4. The percentage of IgA-expressing cells from 3 independent experiments is shown on the right. p-value was determined using two-tailed Student’s t-test; ***p<0,0005.

We focused our analysis on *Fam72a* because of its high-position in the gene-ranking (**Fig. 1B**) and since similarly to AID, its expression is enhanced in wild-type splenic primary B cells undergoing CSR (**Fig. S1D**). *Fam72a* encodes a poorly characterized protein, which has been shown to bind to the nuclear isoform of Uracil DNA Glycosylase (UNG2) in a tryptophan 125-dependent manner^9^ and which is overexpressed in multiple cancers^9^. To validate the screen and determine whether *Fam72a* is required for CSR, we generated *Fam72a^−/−^* CH12 cells using the high-fidelity Cas9 (Cas9-HF1)^17^ (**Fig. S2A and S2B**). Interestingly, we found that the efficiency of CSR was reduced by 50-60 % in four independent *Fam72a^−/−^* CH12 clones when compared to *Fam72a^+/+^* controls (**Fig. 1D**). The CSR defect observed was not due to impaired AID expression (**Fig. S2C**) or defects in proliferation (**Fig. S2D**) or switch region transcription (**Fig. S2E**). We conclude that *Fam72a* is required for efficient CSR in CH12 cells.

To investigate the physiological role of *Fam72a* in CSR and SHM *in vivo*, we obtained a *Fam72a^−/−^* mouse model (**Fig. S3A**) generated by the knockout mouse program (KOMP)^18^. *Fam72a^−/−^* mice gave offspring at Mendelian ratios and showed no obvious deleterious phenotype. *Fam72a^−/−^* mice did not display any specific defects in B cell development, and all B cell populations were found to be represented in normal proportions and numbers in the bone marrow and the spleen (**Fig. 2A, 2B, S3B and S3C**). This suggests that FAM72A is dispensable for RAG1/2- and non-homologous end joining (NHEJ)-dependent V(D)J recombination.

**Figure 2.**
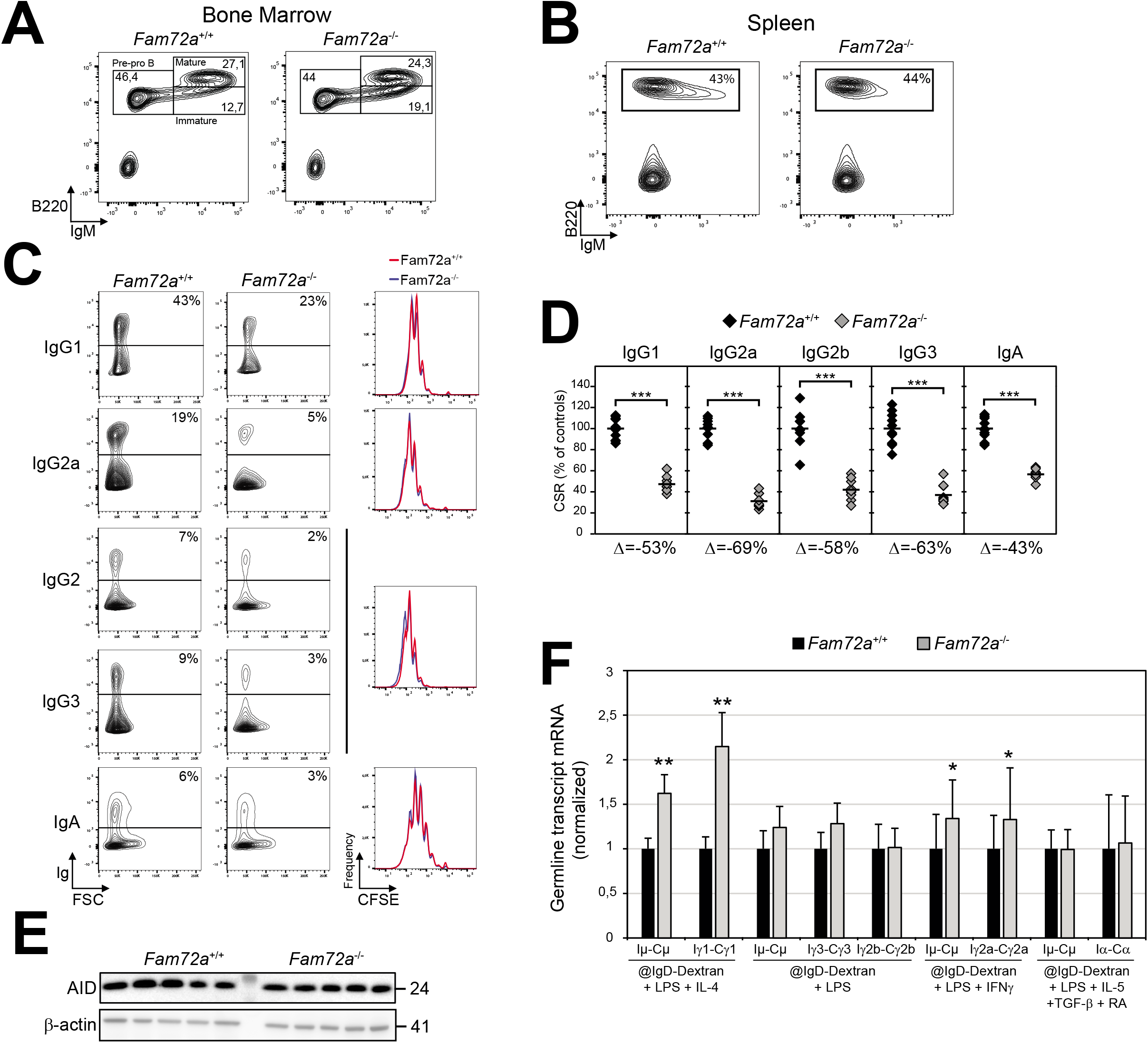
CSR is defective in B cells from Fam72a^−/−^ mice. Flow cytometry analysis of bone marrow **(A)** and spleen **(B)** cells from *Fam72a^+/+^* and *Fam72a^−/−^* mice. Staining antibodies and percentage of cells within gates are indicated. **(C)** Flow cytometry analysis of Ig expression in *Fam72a^+/+^* and *Fam72a^−/−^* splenic B cells cultured for 96h with either LPS + IL-4 + anti-IgD-Dextran (CSR to IgG1), LPS +IFN-γ + anti-IgD-Dextran (CSR to IgG2a) or LPS + anti-IgD-Dextran (CSR to IgG2b and IgG3) or LPS +IL-5 +TGF-β + RA + anti-IgD-Dextran (CSR to IgA). The percentage of Ig-expressing cells is indicated. Representative plots from 2 experiments with 5 mice of each genotype are shown. CFSE dilution analysis is shown on the right. **(D)** Percentage of CSR in *Fam72a^−/−^* B cells relative to *Fam72a^+/+^* B cells for the different isotypes from 10 independent mice is shown on the right. A horizontal line indicates mean values. The difference (Δ) in the percentage of CSR for each isotype between *Fam72a^+/+^* and *Fam72a^−/−^* B cells is indicated below the plot. p-values were determined using two-tailed Student’s t test (***: p<0.0001). **(E)** Western blot analysis for AID and β-Actin in *Fam72a^+/+^* and *Fam72a^−/−^* splenic B cells cultured with LPS, IL-4 and anti-IgD-Dextran for 72h. **(F)** Real-time qPCR analysis for germline transcripts at donor (GLTμ) and acceptor switch regions (GLTγ3, GLTγ1, GLTγ2b, GLTγ2a and GLTα) in *Fam72a^+/+^* and *Fam72a^−/−^* splenic B cells cultured for 96h as in (C). Expression is normalized to Igβ and is presented relative to expression in *Fam72a^+/+^* B cells, set as 1. Mean of 3 independant experiments and the SEM were calculated following the rules for error propagation while calculating a ratio. Statistical analysis was performed using two-tailed Student’s t test (*p < 0,05; **p<0,005).

To determine whether *Fam72a^−/−^* B cells display a defect in CSR, we purified resting splenic B cells, labeled them with CFSE to track proliferation and cultured them under conditions that induce CSR to different isotypes. We found that *Fam72a^−/−^* B cells display a 40-70% reduction in the efficiency of CSR to all isotypes tested (**Fig. 2C and 2D**), which was not due defects in proliferation (**Fig. 2C**), AID expression (**Fig. 2E**), or switch region transcription (**Fig. 2F**). We conclude that deficiency in *Fam72a* results in a B cell intrinsic CSR defect in mouse primary B cells.

To demonstrate that the CSR defect observed is due to deficiency in *Fam72a* and investigate the functional relevance of the FAM72A-UNG2 interaction, we re-expressed FAM72A and the FAM72A^W125R^ or FAM72A^W125A^ UNG2-binding defective mutants^9^ in *Fam72a^−/−^* CH12 B cells. We found that the CSR defect observed was indeed due to the absence of FAM72A, as re-expression of FAM72A (**Fig. S2B**) rescued the CSR defect (**Fig. 3A and 3C**), ruling out unlikely Cas9-HF1 off-target effects. Interestingly, we found that contrary to wildtype FAM72A, the FAM72A^W125R^ or FAM72A^W125A^ mutants were unable to rescue the CSR defect (**Fig. 3A and 3C**). Furthermore, we found that the residual CSR activity observed in *Fam72a*^−/−^ B cells is dependent on the catalytic activity of UNG, as expression of an UNG-inhibitor (Ugi)^19^ abolishes CSR (**Fig. 3B and 3C**). We conclude that the CSR defect observed is due to the lack of FAM72A, that the FAM72A-UNG2 interaction is of functional relevance for CSR and that the residual CSR activity is dependent on the catalytic activity of UNG.

**Figure 3.**
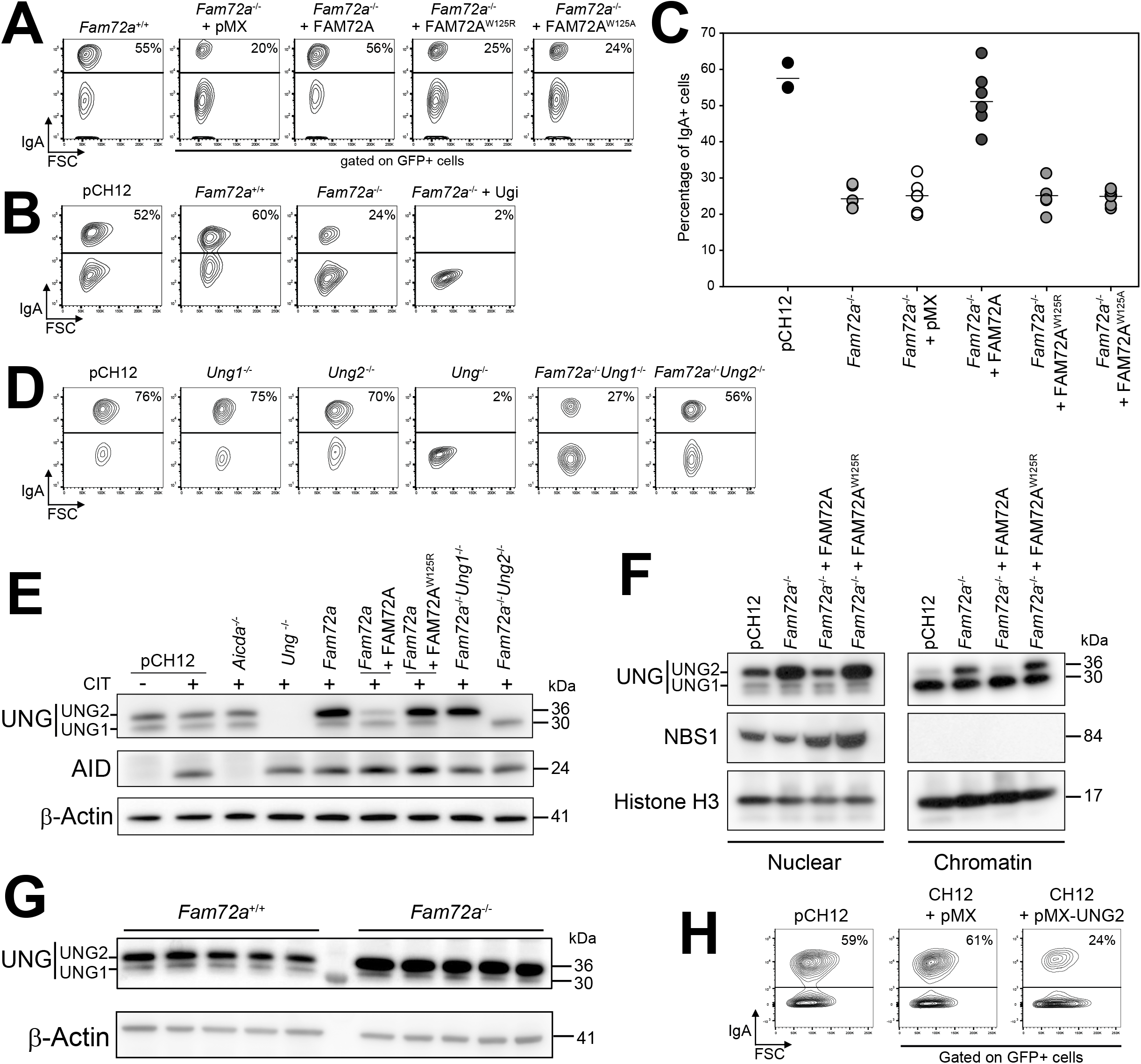
The CSR defect in Fam72a^−/−^ B cells is due to the specific up-regulation of Ung2. **(A)** Flow cytometry analysis of IgA expression in Fam72a^−/−^ CH12 cells transduced with an empty retrovirus (pMX) or expressing FAM72A, FAM72A^W125A^ or FAM72A^W125R^ and cultured for 3 days with TGF-β, anti-CD40 antibody and IL-4. Plots are gated on EGFP expression. The percentage of IgA-expressing cells is indicated. **(B)** Flow cytometry analysis of IgA expression in *Fam72a^+/+^* and *Fam72a^−/−^* CH12 cells expressing (or not) an UNG inhibitor (Ugi) and cultured for 3 days with TGF-β, anti-CD40 antibody and IL-4. The parental cell line (pCH12) was included as a positive control. Representative plots are shown. The percentage of IgA-expressing cells is indicated. **(C)** Plot showing the percentage of IgA-expressing cells from 3 independent experiments. p-value was determined using two-tailed Student’s t-test; ***p<0,0005. **(D)** Flow cytometry analysis of IgA expression in *Fam72a^−/−^, Ung1^−/−^, Ung2^−/−^, Ung^−/−^, Fam72a^−/−^ Ung1^−/−^* and *Fam72a^−/−^ Ung2^−/−^* CH12 cells cultured for 72h with TGF-β, IL-4 and anti-CD40 antibody. The percentage of IgA-expressing cells is indicated. **(E)** Western blot analysis for UNG (UNG1 and UNG2), AID and β-Actin in *Fam72a^−/−^, Ung1^−/−^, Ung2^−/−^ Ung^−/−^ Fam72a^−/−^ Ung1^−/−^, Fam72a^−/−^ Ung2^−/−^* and *Fam72a^−/−^* CH12 cells transduced with a retrovirus expressing FAM72A or FAM72A^W125R^ and cultured for 72h with TGF-β, IL-4 and anti-CD40 antibody (CIT). **(F)** Western blot analysis for UNG (UNG1 and UNG2), NBS1 and Histone H3 on nuclear and chromatin fractions prepared from CH12 cells (pCH12) and *Fam72a^−/−^* B cells expressing FAM72A or FAM72A^W125R^. **(G)** Western blot analysis for UNG (UNG1 and UNG2) and β-Actin protein expression levels in *Fam72a^+/+^* and *Fam72a^−/−^* splenic B cells cultured with LPS, IL-4 and anti-IgD-Dextran for 72h. **(H)** Flow cytometry analysis of IgA expression in CH12 cells transduced with an empty retrovirus or expressing UNG2. Representative plots of three experiments are shown. The percentage of IgA-expressing cells is indicated.

Alternative promoter usage and splicing of the *Ung* gene generates a mitochondrial (UNG1) and a nuclear (UNG2) isoform^20^ (**Fig. S2F**), both of which have been shown to be able to sustain CSR^21^. To examine the relative contribution of both these isoforms relative to FAM72A, we generated *Ung1^−/−^*, *Ung2^−/−^, Ung^−/−^, Fam72a^−/−^ Ung1^−/−^* and *Fam72a^−/−^ Ung2^−/−^* CH12 cell clones using CRISPR/Cas9-HF1 (**Fig. S2F**) and tested their ability to undergo CSR. As shown before^12,21^, while *Ung1^−/−^* and *Ung2^−/−^* cells underwent CSR at wildtype levels (**Fig. 3D and S2G**), CSR in *Ung^−/−^* CH12 cells was completely abolished (**Fig. 3D and S2G**). Surprisingly, while *Fam72a^−/−^ Ung2^−/−^* rescued the CSR defect observed in *Fam72a^−/−^* CH12 cells, *Fam72a^−/−^ Ung1^−/−^* did not (**Fig. 3D and S2G**). This result prompted us to further explore the significance of the FAM72A-UNG2 interaction. For this, we analyzed the expression level of UNG1 and UNG2 in the different single and double knockout CH12 cell lines by

Western blot. Surprisingly, we found that loss-of-function of *Fam72a* results in a significant and specific up-regulation of the UNG2 isoform (**Fig. 3E**), which accumulated on chromatin (**Fig. 3F**). This suggests that FAM72A controls the protein level of UNG2 and its access to chromatin. This phenotype, which was not at the mRNA level (**Fig. S2H**), was also observed in *Fam72a^−/−^* primary B cells (**Fig. 3G**). Furthermore, when we overexpressed FAM72A in *Fam72a^−/−^* CH12 cells (**Fig. S2B**), the up-regulation of UNG2 was almost abolished (**Fig. 3E**). Significantly, overexpression of the FAM72A^W125A^ mutant (**Fig. S2B**), which does not interact with UNG2^9^, did not down-regulate UNG2 (**Fig. 3E**). We conclude that FAM72A specifically regulates the protein level of the UNG2 isoform by controlling its degradation and that the CSR defect observed in *Fam72a^−/−^* and *Fam72a^−/−^ Ung1^−/−^* cells is due to increased levels of the UNG2 isoform. Consistent with this, we found that overexpression of UNG2 in wildtype CH12 cells was sufficient to suppress CSR (**Fig. 3H**). Therefore, it appears that the maintenance of the physiological levels of UNG2 is of critical importance for the efficiency of CSR and explains why mutants, with defective catalytic activity of UNG2, reconstitute CSR better than wildtype UNG2 in *Ung*^−/−^ B cells^22–24^. Our results suggest that FAM72A, by controlling the level of UNG2, influences the usage of error-prone *versus* error-free DNA repair in response to AID-induced uracils in DNA.

Activated B cells expressing AID display mutations at switch regions during CSR^25^ and at a region spanning immunoglobulin variable regions during SHM^3^. To determine whether deficiency in *Fam72a* and the concomitant up-regulation of UNG2 have an influence in the frequency and pattern of AID-induced mutations, we sequenced the 5’ end of the Sμ switch regions in *Fam72a^−/−^* and control CH12 cells that were stimulated to undergo CSR for three days. Interestingly, we found that the mutation frequency was reduced by 50% in *Fam72a^−/−^* CH12 cells when compared to controls (**Fig. S4A**). To confirm whether deficiency in *Fam72a* results in lower AID-induced mutation frequency, we analyzed the J_H_4 intron (J_H_4i), a sequence which gets heavily mutated by AID during SHM *in vivo*^26^, in germinal center B cells isolated from the Peyer’s patches of unimmunized *Fam72a^+/+^* and *Fam72a^−/−^* mice (**Fig. 4A**). Interestingly, we found that the mutation frequency at the J_H_4i sequence was drastically reduced (five-fold) in *Fam72a^−/−^* B cells, when compared to controls (**Fig. 4B, 4C and S4B**). Consistent with UNG2 up-regulation, we found that the percentage of transition mutations at C:G base pairs, which are generated by replication over U:G mismatches were reduced from 57 % in *Fam72a*^+/+^ to 16 % in *Fam72a*^−/−^ B cells (**Fig. 4C and S4B**). Conversely, transversion mutations at C:G base pairs, which are generated through replication over UNG-generated abasic sites, increased from 43 % in *Fam72a^+/+^* to 84 % in *Fam72a^−/−^* B cells (**Fig. 4C** and **S4B**). This phenotype is diametrically opposed to *Ung^−/−^* B cells^12^, in which transition mutations at C:G base pairs reaches 95 % and transversion mutations at C:G base pairs are suppressed to 5 %^12^. These results support the hypothesis that in *Fam72a*-deficient B cells, where UNG2 is upregulated, AID-induced uracils are more efficiently excised from DNA by UNG2 and that error-free repair is enforced. Furthermore, enhanced uracil excision would result in fewer U:G mismatches, rendering MMR less effective in generating mutations at A:T base pairs surrounding the deaminated cytosines. Therefore, the level of UNG2 needs to be tightly controlled to trigger error-prone DNA repair, which is essential for antibody diversification through SHM and CSR.

**Figure 4.**
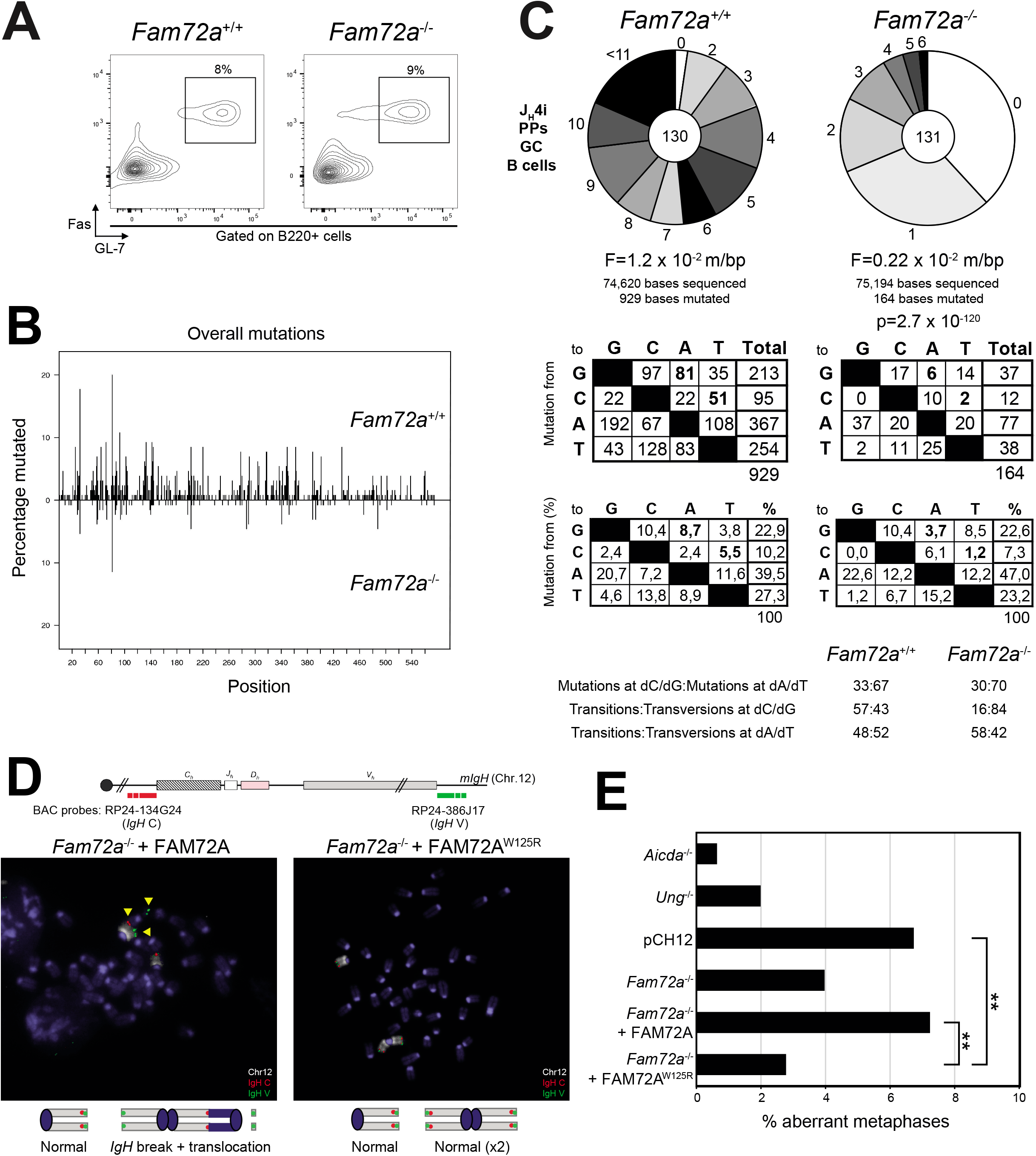
Deficiency in Fam72a shifts the balance toward error-free DNA repair. **(A)** Flow cytometry analysis of germinal center B cells (B220+ Fas+ GL-7+) isolated from the Peyer’s patches of unimmunized *Fam72a^+/+^* and *Fam72a^−/−^* mice. Plots are gated on B220+ cells. **(B)** Distribution of mutations at the J_H_4 intron (J_H_4i) in *Fam72a^+/+^* (top) and *Fam72a^−/−^* (bottom) sequences obtained from germinal center B cells (B220+ Fas+ GL-7+) isolated from the Peyer’s patches (PPs) of unimmunized *Fam72a^+/+^* and *Fam72a^−/−^* mice. **(C)** Pie charts depict the proportion of J_H_4i sequences with the indicated number of mutations. The total number of sequences analyzed is indicated in the center and the mutation frequency below. Tables depicting the mutation profiles are shown below. Statistical significance was determined using the SHMtool server. **(D)** Upper panel: Schematic representation of the *Igh* locus with positions of the BACs used for generation of DNA FISH probes. Lower panel: DNA-FISH on representative metaphases from day 2 stimulated *Fam72a^−/−^* cells complemented with FAM72A or FAM72A^W125R^ using the Igh C BAC probe (red) combined with Igh V BAC probe (green) and chromosome 12 paint (white). Yellow arrowheads point to broken or translocated chromosome 12. **(E)** Percentage of aberrant metaphases from day 2 stimulated cells of the indicated genotype harboring *Igh* locus aberrations. See also Table S1. ** p<0.01.

CSR is sensitive to the expression level of AID and concomitantly to the amount of AID-induced DSBs, which are obligatory intermediates^3,10^. Indeed, AID is haplo-insufficient and CSR is reduced in AID^+/-^ B cells^27^. The CSR defect observed in *Fam72a^−/−^* B cells could be explained by enforced error-free repair due to the up-regulation of UNG2, which in turn would lead to the generation of fewer DSBs at switch regions. In wildtype B cells, a small fraction of AID-induced DSBs can be detected in the form of chromosome breaks or translocations^28^. To determine whether up-regulation of UNG2 in the absence of FAM72A might influence the level of *IgH* DSBs, we conducted fluorescence in situ hybridization experiments using two *IgH*-specific probes and chromosome 12 paint in activated *AID^−/−^*, *Ung*^−/−^, *Fam72a^+/+^* and *Fam72a^−/−^* cells reconstituted (or not) with FAM72A or FAM72A^W125R^ and quantified aberrant metaphases in these cells (**Fig. 4D, 4E and Table S1**). We found lower levels of aberrant metaphases in *Fam72a^−/−^* cells or *Fam72a^−/−^* cells reconstituted with FAM72A^W125R^ as compared to wild type and *Fam72a^−/−^* cells reconstituted with wild type FAM72A, suggesting that FAM72A-mediated UNG2 upregulation controls the level of AID-induced DSBs during CSR. We conclude that the CSR defect observed in the absence of *Fam72a* is due to the inefficient generation of AID-induced DSBs.

Our results are consistent with a model in which FAM72A interacts with UNG2 to control its physiological level by triggering its degradation. Consequently, deficiency in *Fam72a* leads to the specific upregulation of the UNG2 isoform and its accumulation on chromatin. This is sufficient to enforce uracil excision, resulting in a reduction in the efficiency of SHM and CSR. It is possible that supraphysiological levels of UNG2 in the absence of FAM72A tilt the balance toward error-free BER by favoring the recruitment of polymerase β^5,6,29,30^. A shift in the balance from error-free to error-prone DNA repair in response to uracils in DNA has significant implications in the onset of cancer. As *Fam72a* is overexpressed in several types of tumors and transformed cell lines^9^ and as *Ung^−/−^* mice develop B cell lymphomas^31–33^, it is possible that *Fam72a* overexpression would suppress the levels of UNG2 leading to inefficient uracil excision, enforcing error-prone DNA repair and making these cells more susceptible to accumulate mutations and hence more prone to tumorigenesis.

## Acknowledgements

B. Kavli for the anti-UNG1/2 antibody; L. Brino for help with gRNA library sub-cloning and CRISPR/Cas9 screen analysis; B. Jost, C. Thibault-Carpentier, D. Plassard and D. Dembele for gRNA high-throughput sequencing; B. Heller for cell culture; C. Ebel and M. Philipps for cell-sorting; M. Selloum, D. Ali-Hadji and I. Gonçalves Da Cruz for animal care. J. Moritz was supported by the Ministère de l’Enseignement Supérieur et de la Recherche, France and the Fondation ARC. M. Rogier was supported by La Ligue Contre le Cancer. This study was supported by grants from the Fondation Recherche Médicale (Equipe FRM EQU201903007818), Institut National du Cancer (INCa_13852), Fondation ARC (ARCPJA32020060002061) and by the grant ANR-10-LABX-0030-INRT, a French State fund managed by the Agence Nationale de la Recherche under the program Investissements d’Avenir labelled ANR-10-IDEX-0002-02.

## Author Contributions

B.R.S.M. conceived the project. M.R., J.M., I.R., C.L., A-S.T.C-P., A.A. and V.H. performed experiments and analyzed the data. M.R., J.M. and B.R.SM. wrote the manuscript. L.D. and B.R.S.M directed overall research and edited the manuscript.

Supplementary Information is available for this paper.

Correspondence and requests for materials should be addressed to B.R.S.M.

## Methods

### CH12^Cas9^ cell generation

CH12 cells were transduced with retroviral supernatants obtained by transfecting Bosc23 cells with a retrovirus (pMX-26) expressing a Cas9-mCherry-P2A-Hygromycin cassette (**Fig. S1A**). After 72h of hygromycin selection (Sigma; 300 μg/mL), mCherry+ cells were sorted into 96-well plates using a cell sorter FACS ARIA Fusion (Becton Dickinson) and cultured for 10 days with hygromycin (Sigma; 300 μg/mL). Individual clones were then tested for their ability to express Cas9 by western-blot (**Fig. S1A)**, and their ability to undergo efficient CSR after stimulation (**Fig. S1A)**. To test the functionality of Cas9, clones were transfected with gRNAs known to be effective (see Table S2 for gRNA sequences). Three days later, PCR was performed to identify clones with Cas9-mediated deletions (**Fig. S1A)**. Clone #11, which had the highest Cas9 activity and which displayed a CSR efficiency similar to the parental CH12 cell line was selected to conduct the screen.

### Sub-cloning of the mBrie gRNA library

The mouse Brie CRISPR knockout pooled library (lentiCRISPRv2 backbone; a gift from David Root and John Doench (Addgene #73633) was sub-cloned into the pMX-28 retroviral vector (**Fig. S1B**) as described^16^ to generate the pMX-mBrie gRNA library. Briefly, the U6p-gRNA-scaffold cassette was amplified by PCR (see Table S2 for primers), digested with BamHI and a NotI restriction enzymes and ligated into pMX-28 (**Fig. S1B**). Stbl4 electrocompetent cells (ThermoFisher) were electroporated using Eporator Eppendorf (Program1-1700V), plated on ampicillin agar plates and incubated at 37°C for 15 hours. Plasmid DNA was extracted and the U6-gRNA-scaffold cassette was amplified by PCR to add sequencing adaptors and barcodes (see Table S2 for primers) and then analyzed by deep sequencing to determine gRNA representation and library uniformity as described^16^.

### CRISPR/Cas9 knockout screen

CH12^Cas9^ cells were transduced with the pMX-mBrie gRNA library in quadruplicate at a multiplicity of infection (MOI) of 0.3 and with a 300X coverage, as previously described^16^. Transduced cells were selected with puromycin (Sigma; 0.5 μg/mL) and hygromycin (Sigma; 300 μg/mL) for 15 days. Selected cells were then induced to undergo CSR with TGF-β (1 ng/ml; R&D Systems Europe), IL-4 (5 ng/ml; Peprotech) and an anti-CD40 antibody (200 ng/ml; eBioscience) for 72h. IgM+ and IgA+ cells were sorted using anti-IgM and anti-IgA coupled magnetic beads (Mitentyi). Population purity was assessed by flow cytometry (**Fig. S1C)** using a Fortessa flow cytometer (Becton Dickinson). Genomic DNA from 24 million of cells was extracted using phenol/chloroform and subjected to PCR (1x (95°C for 2min)/ 28-31X (95°C for 15s, 65°C for 20s, 72°C for 30s) and 72°C for 3 min) to amplify gRNA sequences using staggered primers having Illumina adaptors and barcodes (see Table S2 for primers). Multiplexed samples were submitted to high-throughput sequencing (1×50 bp) on an Illumina HiSeq4000 sequencer at the GenomEast sequencing platform of IGBMC. gRNA counts were extracted from raw data using the PoolQ software from the Broad Institute. gRNA counts were normalized according as described^16^. Ranking of candidate genes was performed using MaGeCK^34^. Candidate genes (p<0.05) and having more than 2 effective gRNAS were selected.

### Cell culture

Bosc23 cells were cultured in DMEM supplemented with glucose (4,5g/L), 10% of heat-inactivated fetal calf serum, penicillin-streptomycin (100U/mL) and sodium pyruvate (1mM). CH12 cells and primary B cells were cultured in RPMI supplemented with 10% of heat-inactivated fetal calf serum, HEPES (10 mM), penicillin-streptomycin (100 U/mL), Sodium Pyruvate (1 mM) and β-mercaptoethanol (50 μM).

### CSR assays

CH12 cells were cultured for 72h in the presence of TGF-β (1 ng/ml; R&D Systems Europe), IL-4 (5 ng/ml; Peprotech) and an anti-CD40 antibody (200 ng/ml; eBioscience). Cells were then stained with an anti-IgA-PE antibody (Southern Biotech) to assess CSR by flow cytometry. Prior to analysis, DAPI was added to discriminate dead cells. Samples were analyzed using a Fortessa flow cytometer (Becton Dickinson) and the FlowJo software.

### Primary B cell cultures

Splenic resting B cells were purified using anti-CD43 magnetic beads (Miltentyi), labeled with CFSE (COMPANY) and cultured from 4 days with a combination of LPS (25 μg/ml, Sigma-Aldrich), IL-4 (25 μg/ml, Preprotech), anti-IgD-Dextran (6 ng/ml, Fina Biosolutions), IFN-γ (100 ng/ml, Preprotech), IL-5 (5 ng/ml, R&D Systems), TGF-β (3 ng/ml, R&D Systems) and Retinoic Acid (RA; 0.3 ng/ml, Sigma-Aldrich).

### RTq-PCR

RNA and cDNA were prepared using standard techniques. qPCR was performed in triplicates using Roche LightCycler 480 Probes Master mix Universal Probe Library (UPL) in combination with appropriate UPL probes (**Table S2**). Transcript quantities were calculated relative to standard curves and normalized to HPRT or Igβ mRNA.

### Retroviral transductions

CH12 cells were transduced with retroviral supernatants obtained by transfecting Bosc23 cells with an empty retrovirus (pMX-PIE; Puromycin-IRES-EGFP) or expressing mFam72a (pMX-Fam72a), o mFam72a^W125R^ (pMX-mFam72a^W125R^), mFam72a^W125A^ (pMX-Fam72a^W125A^), mUng2 (pMX-mUng2) or Ugi (pMX-Ugi). Transduced cells were then selected with puromycin (1 μg/ml) for 7 days and submitted to CSR assays.

### Generation of CH12 knockout clones

CH12 cells were transfected by electroporation using the Neon transfection System (ThermoFisher) with a plasmid expressing one or two gRNAs targeting a critical exon (see Table S2 for gRNA sequences) and co-expressing the high-fidelity Cas9 nuclease^17^ coupled to EGFP. 24h after transfection, individual EGFP-positive cells were sorted into 96-well plates using a cell sorter FACS ARIA Fusion (Becton Dickinson) and cultured for 10 days. Clones were then genotyped by PCR and sequencing.

### Sμ and JH4i somatic hypermutation analysis

For Sμ mutation analysis, genomic DNA was extracted from CH12 cells cultured for 3 days with TGF-β (1 ng/ml; R&D Systems Europe), IL-4 (5 ng/ml; Peprotech) and an anti-CD40 antibody (200 ng/ml; eBioscience). For JH4i mutation analysis, genomic DNA was extracted from B cells isolated from the Peyer’s patches and sorted by flow cytometry using anti-B220-PE-Cy7 (eBiosciences), anti-Fas-PE (BD Pharmingen) and anti-GL7-Pacific Blue (BioLegend) antibodies. Sμ and JH4i sequences were amplified by PCR (1x 98°C for 30s, 35x (98°C for 10s, 70°C for 10s and 72°C for 30s) and 1x 72°C for 5 min) using the Q5 polymerase (New England BioLabs) and cloned into pUC57 using the MEGAWHOP^35^ method. Inserts were sequenced (Sanger sequencing) using the M13 Forward universal primer. Sequences were aligned with Lasergene (DNASTAR) and analyzed with the SHMTool^36^ server.

### Cloning

All gRNA/Cas9HF1 plasmids were generated through golden gate cloning^37^. cDNAs were cloned into the pMX-PIE retrovirus through SLICE^38^.

### Mice

*Fam72a*^−/−^ mice (C57BL6) were generated by the knockout mouse program (KOMP)^18^. Mice were bred under pathogen-free (SPF) conditions. In all experiments 8-12 week old age-matched littermates were used. Animal work was performed under protocols approved by an ethics committee (APAFIS#23104-2019112915476749).

### IgH FISH

Metaphases were prepared using standard procedures^39^. DNA FISH on metaphases spreads was performed as previously described^39^ using BAC probes RP24-134G24 (5’ Igh C) and RP24-386J17 (3’ Igh V) and XCyting Mouse Chromosome 12 (Orange) paint from MetaSystems. Metaphases were imaged using a ZEISS AxioImager.Z2 microscope and the Metafer automated capture system (MetaSystems), and counted manually.

### Western blot analysis

Proteins extracts were prepared using standard techniques. Proteins were separated by SDS-PAGE using gradient gels (4-12%; Invitrogen), transferred to PVDF membranes (Immobilon; Millipore) and analyzed using anti-AID^40^, anti-β-Actin (Sigma), anti-UNG1/2 (Gift from B. Kavli) antibodies. Cell fractionation experiments were performed as previously described^41^.

### B cell development

Total splenic or bone marrow cells were labeled with suitable antibodies and analyzed on an FACS-Fortessa flow cytometer (Becton Dickinson).

---

**Figure S1.**
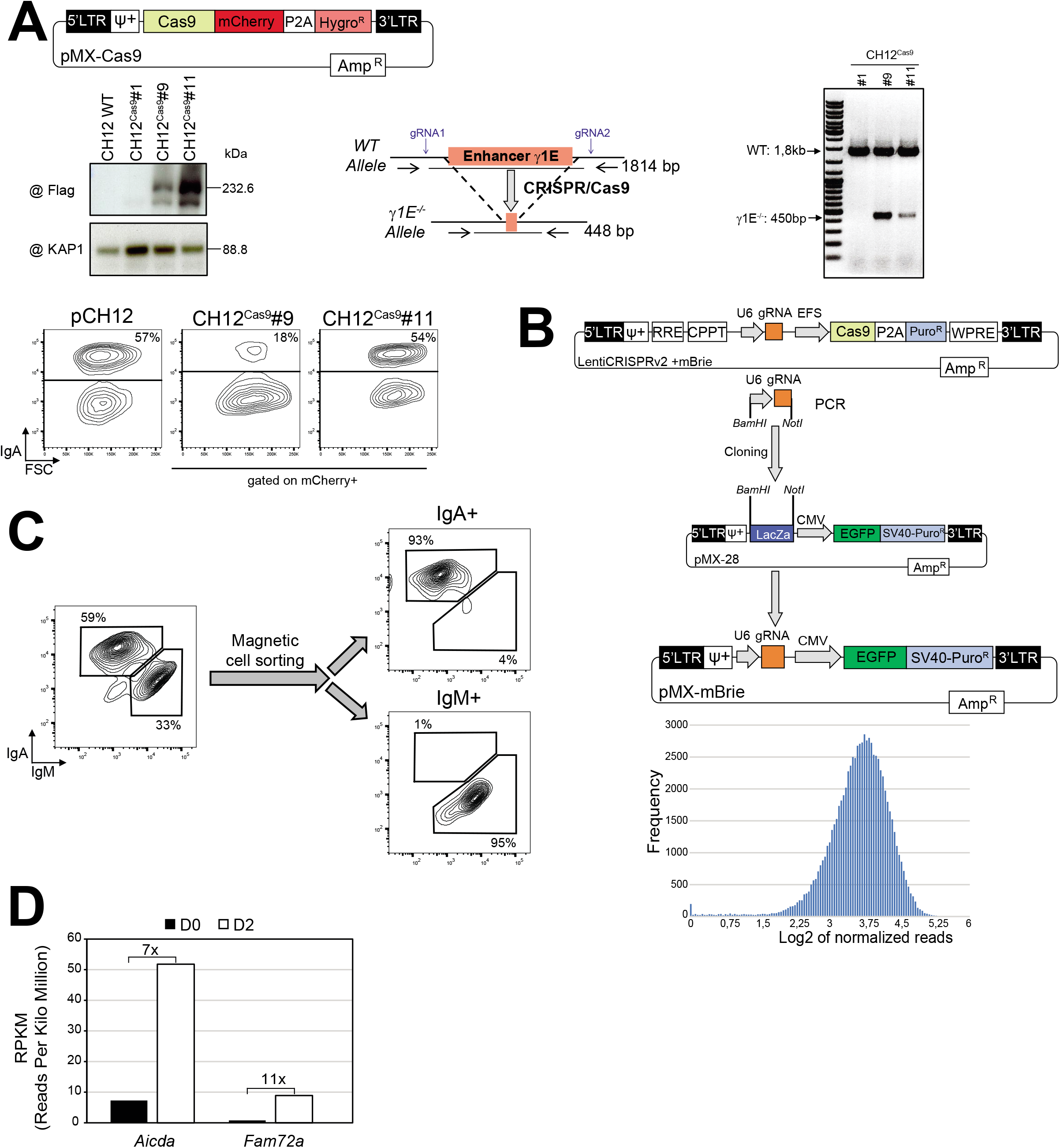
**(A)** Generation and validation of Cas9-expressing (CH12Cas9) cells. Clones were verified by Western blot for Cas9 expression, by PCR for Cas9-induced deletion of the enhancer γ1 and by flow cytometry for CSR. Clone #11 was chosen for the screen. **(B)** The U6-gRNA sequence was amplified by PCR from the mBrie library and subcloned into pMX-28 using BamHI and NotI restriction enzymes to generate the pMX-mBrie gRNA library. gRNA representation was analyzed by high-throughput sequencing. **(C)** Purity of IgM+ and IgA+ sorted populations was verified by flow cytometry. **(D)** Number of reads at the *Aicda* and *Fam72a* genes in wildtype splenic B cells before and after 48h in culture with LPS and IL-4, as determined by mRNA-Seq.

**Figure S2.**
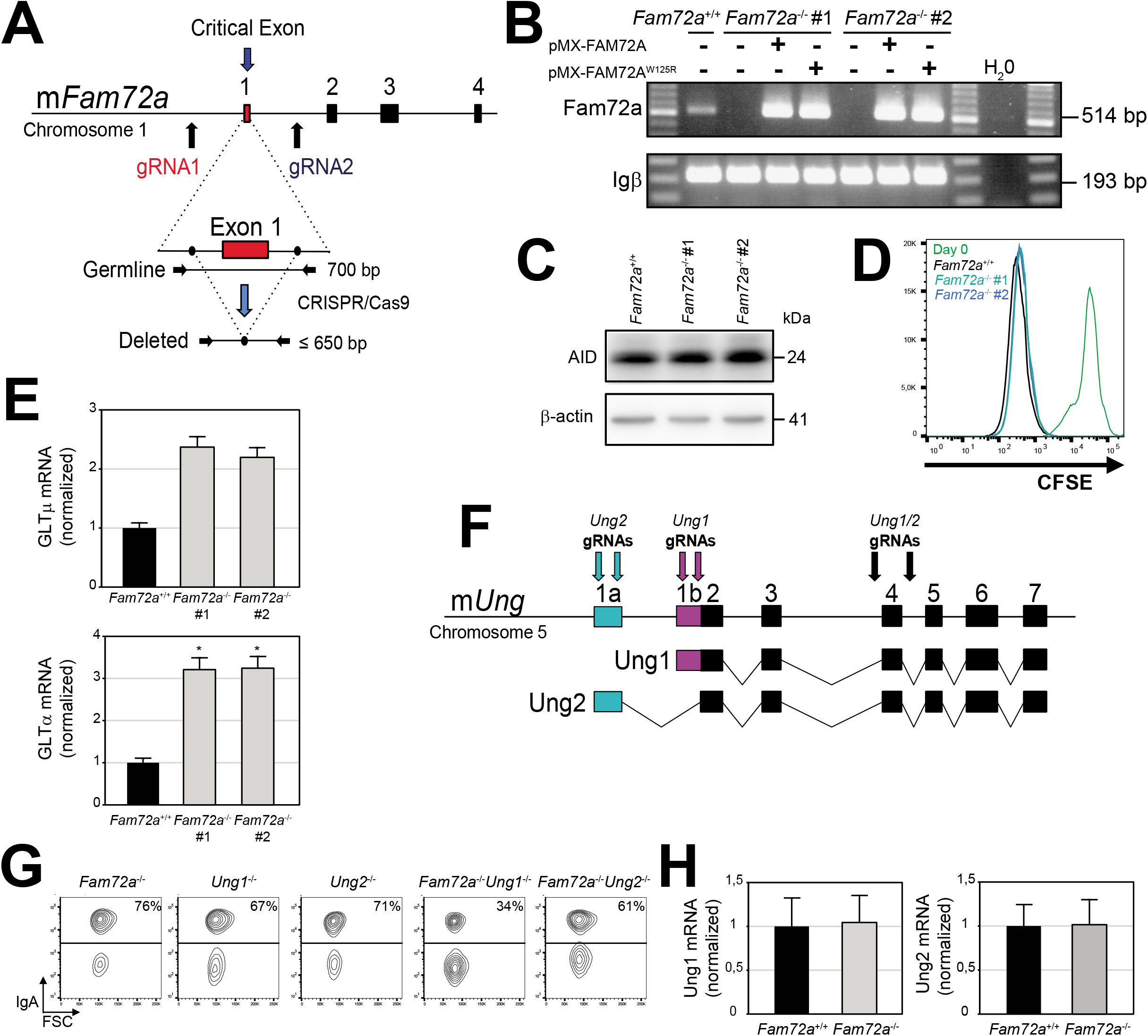
**(A)** Schematic representation of the murine *Fam72a* locus and location of the gRNAs used to generate *Fam72a^−/−^* clones using CRISPR/Cas9-HF1. **(B)** RT-PCR analysis for Fam72a and Igβ in *Fam72a^+/+^* and *Fam72a^−/−^* CH12 cells transduced (or not) with a retrovirus expressing FAM72A, FAM72A^W125R^ or FAM72A^W125A^. **(C)** Western blot analysis for AID and β-actin in *Fam72a^+/+^* and two independent *Fam72a^−/−^* CH12 cell clones cultured for 3 days with TGF-β, IL-4 and anti-CD40 antibody. **(D)** CFSE dye-dilution analysis by flow cytometry in *Fam72a^+/+^* and two independent *Fam72a^−/−^* CH12 cell clones cultured for 3 days with TGF-β, IL-4 and anti-CD40 antibody. **(E)** RT-qPCR analysis for GLTμ and GLTα in *Fam72a^+/+^* and *Fam72a^−/−^* CH12 cells cultured for 3 days with TGF-β, IL-4 and anti-CD40 antibody. Triplicates were normalized to the abundance of Igβ and are expressed relative to *Fam72a^+/+^* controls. Statistical significance was determined by a two-tailed Student’s t-test (*p-value<0,05). Data are representative of 3 experiments.**(F)** Schematic representation of the murine *Ung* locus and location of the gRNAs targeting *Ung2* (exon 1a; blue), *Ung1* (exon 1b; purple) or *Ung* (exon4; black) used to generate *Ung1^−/−^, Ung2^−/−^, Ung^−/−^, Fam72a^−/−^ Ung1^−/−^* and *Fam72a^−/−^ Ung2^−/−^* CH12 cell clones using CRISPR/Cas9-HF1. **(G)** Flow cytometry analysis of IgA expression in additional *Ung1^−/−^, Ung2^−/−^, Fam72a^−/−^ Ung1^−/−^* and *Fam72a^−/−^ Ung2^−/−^* CH12 cell clones cultured for 3 days with TGF-β, IL-4 and anti-CD40 antibody. **(H)** RT-qPCR analysis for Ung1, Ung2 and Igβ in *Fam72a^+/+^* and *Fam72a^−/−^* splenic B cells cultured for 4 days with LPS, IL-4 and anti-IgD-Dextran. Triplicates were normalized to the abundance of Igβ and are expressed relative to *Fam72a^+/+^* controls.

**Figure S3.**
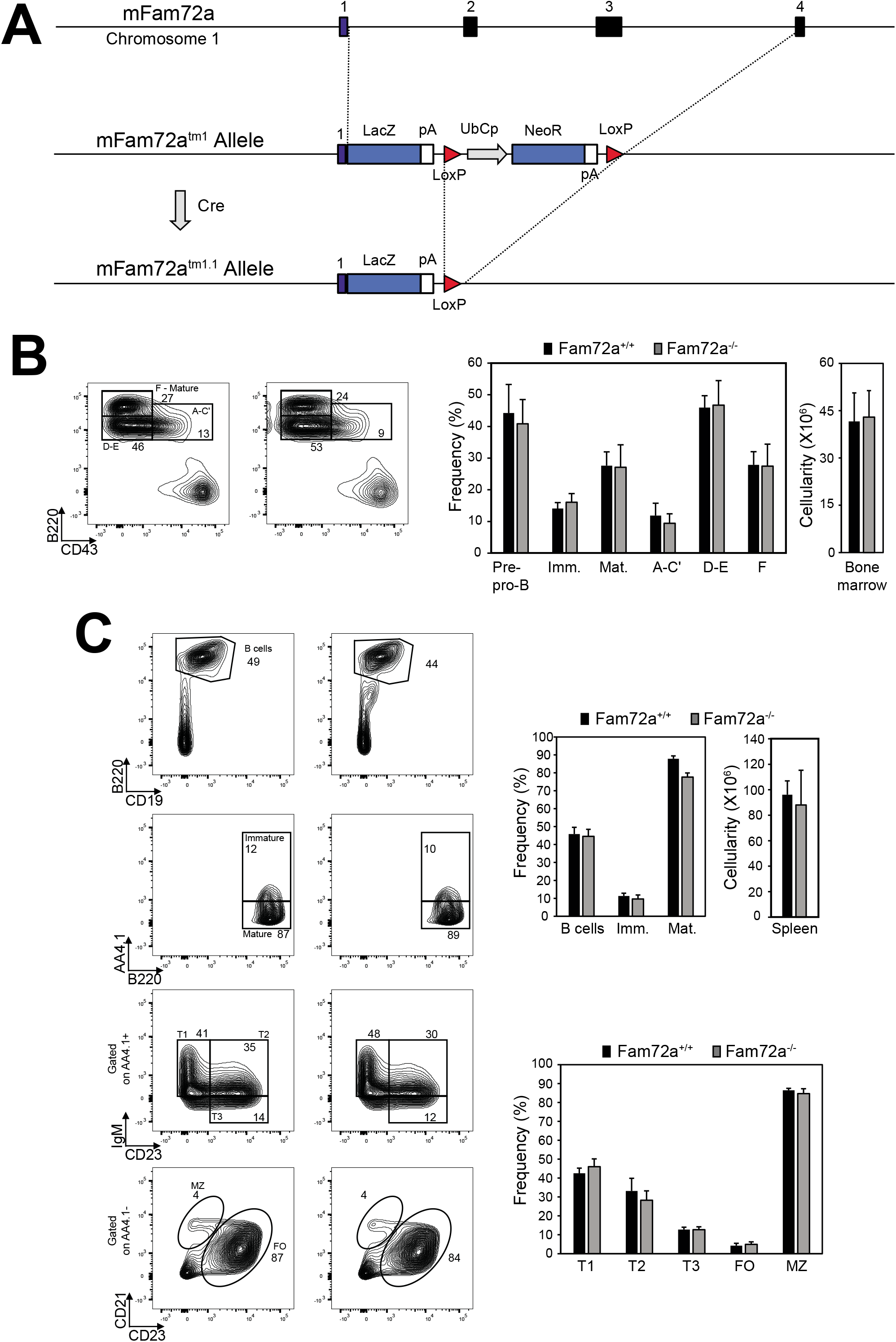
**(A)** Schematic representation of the murine *Fam72a* locus and strategy for the generation of *Fam72a^−/−^* mouse model. Flow cytometry and cellular analysis of B cell populations in the bone marrow **(B)** or in the spleen **(C)** from *Fam72a^+/+^* and *Fam72a^−/−^* mice, using the indicated cell surface markers. The data are representative of 5 mice per genotype.

**Figure S4.**
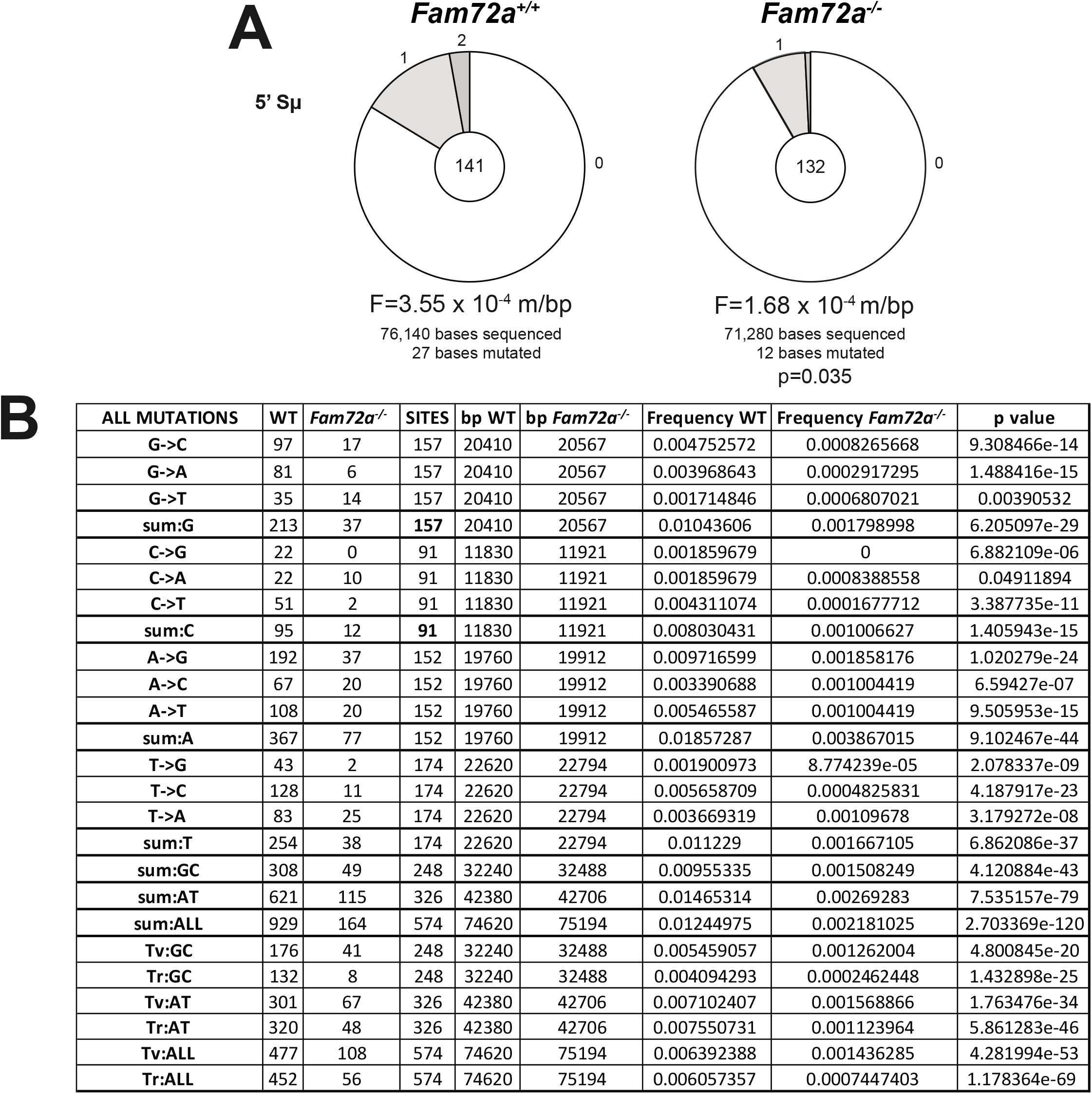
**(A)** Sμ mutation analysis in *Fam72a^+/+^* and *Fam72a^−/−^* CH12 cells cultured for 3 days with TGF-β, IL-4 and anti-CD40 antibody. Pie charts depict the proportion of Sμ sequences with the indicated number of mutations. The mutation frequency per base pair sequenced is shown below. The total number of sequences analyzed is indicated in the center. Statistical significance was determined with the Student’s t test. **(B)** Mutation analysis at J_H_4i sequences performed with the SHMTool server.

**Table S1.** IgH FISH analysis.

| Genotype | Expmt # | Metaphase<br>s with *Igh*<br>breaks | Metaphases<br>with *Igh*<br>translocation<br>s | Total<br>aberrant<br>metaphase<br>s | Total<br>normal<br>metaphase<br>s | Total<br>metaphase<br>s analyzed | %<br>abnormal<br>metaphase<br>s |
| --- | --- | --- | --- | --- | --- | --- | --- |
| *Aicda*<sup>-/-</sup> | 1 | 2 | 0 | 2 | 248 | 250 | 0,8 |
|  | 2 | 1 | 0 | 1 | 219 | 220 | 0,5 |
|  | **TOTAL** | **3** | **0** | **3** | **467** | **470** | **0,6** |
| *Ung*<sup>-/-</sup> | 1 | 4 | 0 | 4 | 235 | 239 | 1,7 |
|  | 2 | 4 | 1 | 5 | 209 | 213 | 2,3 |
|  | **TOTAL** | **8** | **1** | **9** | **444** | **452** | **2,0** |
| pCH12 | 1 | 11 | 6 | 16 | 233 | 249 | 6,4 |
|  | 2 | 17 | 2 | 19 | 252 | 271 | 7,0 |
|  | **TOTAL** | **28** | **8** | **35** | **485** | **520** | **6,7** |
| *Fam72a*<sup>-/-</sup> | 1 | 6 | 4 | 10 | 242 | 252 | 4,0 |
|  | **TOTAL** | **6** | **4** | **10** | **242** | **252** | **4,0** |
| *Fam72a*<sup>-/-</sup><br>+FAM72A | 1 | 10 | 3 | 13 | 241 | 254 | 5,1 |
|  | 2 | 15 | 0 | 15 | 118 | 133 | 11,3 |
|  | **TOTAL** | **25** | **3** | **28** | **359** | **387** | **7,2** |
| *Fam72a*<sup>-/-</sup><br>+FAM72A<sup>W125R</sup> | 1 | 3 | 0 | 3 | 157 | 160 | 1,9 |
|  | 2 | 7 | 1 | 8 | 228 | 236 | 3,4 |
|  | **TOTAL** | **10** | **1** | **11** | **385** | **396** | **2,8** |

| Statistical analysis (Fisher exact test) |  |  | p value | Level |
| --- | --- | --- | --- | --- |
| *Fam72a*<sup>+/+</sup> | vs | *Fam72a*<sup>-/-</sup> | 0.14188 | ns |
| *Fam72a*<sup>+/+</sup> | vs | *Fam72a*<sup>-/-</sup> + FAM72A | 0.79264 | ns |
| *Fam72a*<sup>+/+</sup> | vs | *Fam72a*<sup>-/-</sup> + FAM72A<sup>W125R</sup> | 0.00873 | ** |
| *Fam72a*<sup>-/-</sup> | vs | *Fam72a*<sup>-/-</sup> + FAM72A | 0.12234 | ns |
| *Fam72a*<sup>-/-</sup> | vs | *Fam72a*<sup>-/-</sup> + FAM72A<sup>W125R</sup> | 0.49575 | ns |
| *Fam72a*<sup>-/-</sup> + FAM72A | vs | *Fam72a*<sup>-/-</sup> + FAM72A<sup>W125R</sup> | 0.00481 | ** |

**Table S2.**
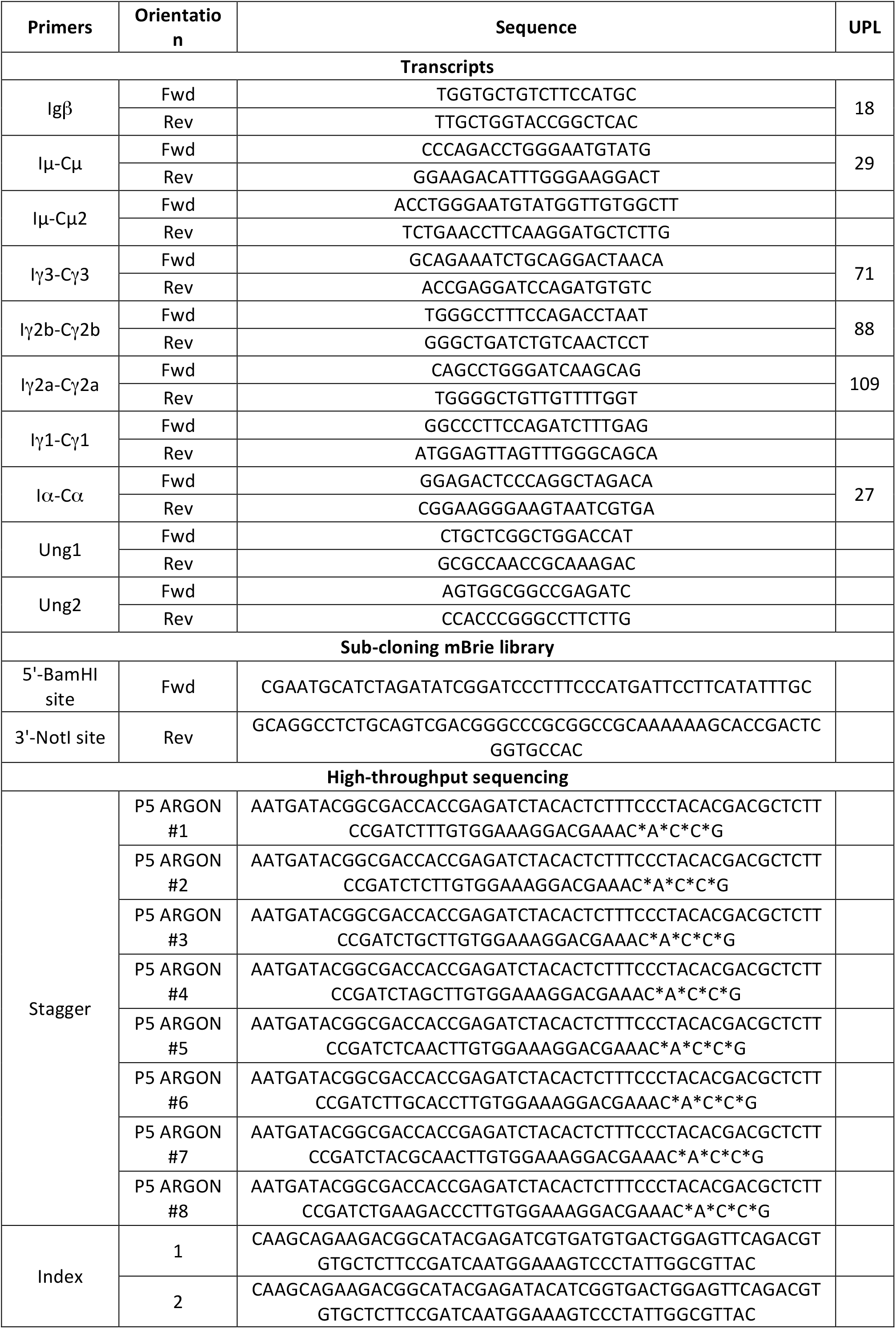
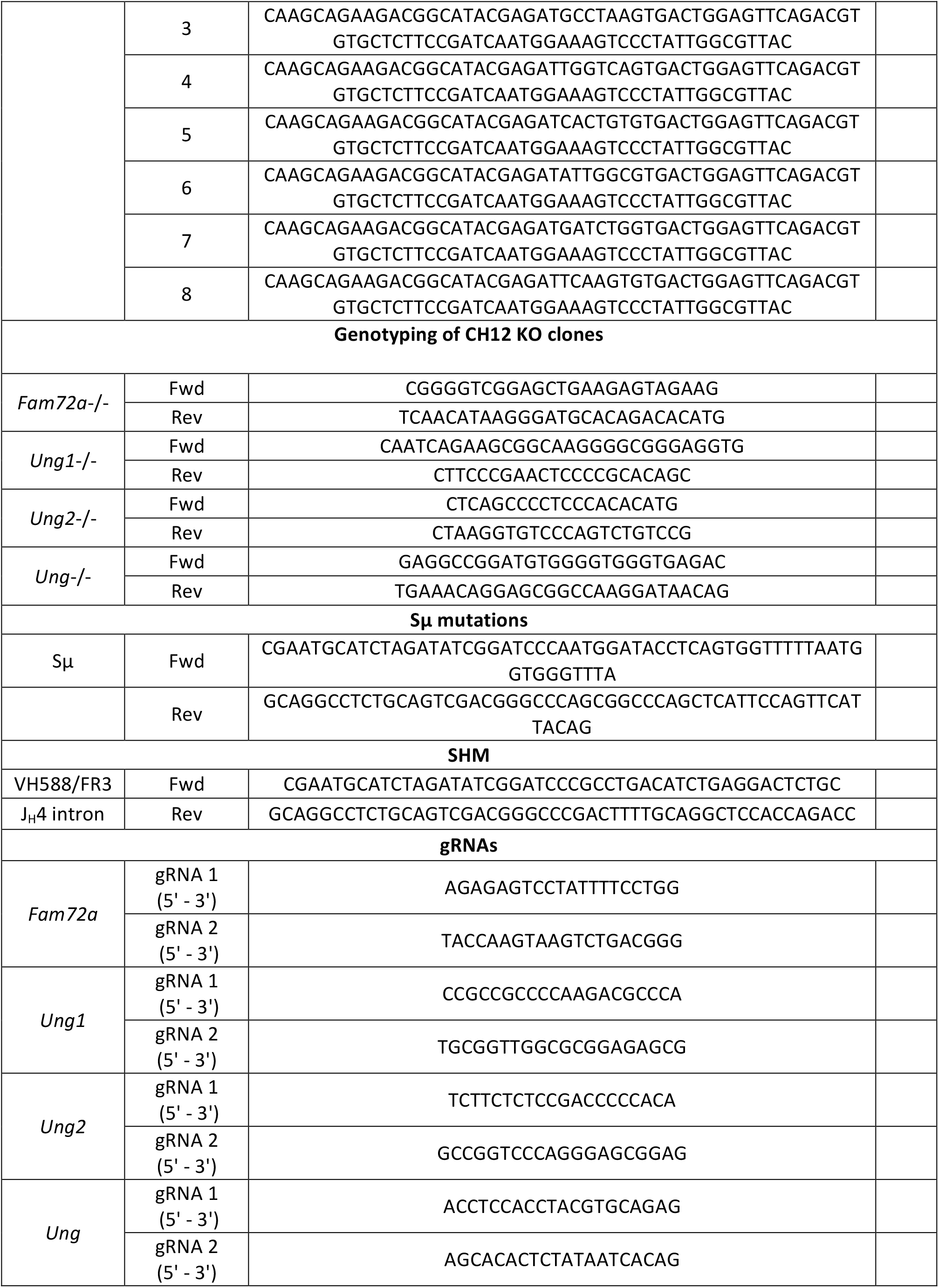
Primers and gRNAs.

## Notes

### Competing Interest Statement

The authors have declared no competing interest.

